# Interaction between task-evoked and spontaneous brain activities: Insights from two event-related fMRI studies

**DOI:** 10.1101/2023.09.16.557985

**Authors:** Siddharth Nayak, Seyed Hani Hojjati, Sindy Ozoria-Blake, Farnia Feiz, Jacob Shteingart, Antonio Fernandez Guerrero, Qolamreza R Razlighi

## Abstract

Human brain activities measured by fMRI can be categorized into at least two distinct categories; 1) spontaneous: slow (< 0.1 Hz) but inter-regionally coherent fluctuations in MR signal, and 2) task-evoked: change in the MR signal due to external stimulation. To date, there is no consensus in the field about the roles of spontaneous brain activity and whether it interacts with task-evoked activity. This is mainly because direct measurements of spontaneous activity during task performance is challenging, if not impossible. Here for the first time, we used a data-driven approach to obtain the time course of the spontaneous brain activity in two of the well-studied brain networks; 1) the Default mode network (DMN), where the strongest task-evoked negative BOLD response is also observed and 2) the dorsal attention network (DAN), where some of the strongest task-evoked positive BOLD response is observed. Voxel-wise multiple regression analysis is used to model fMRI timeseries with anticipated task-evoked BOLD response, the obtain time course of the spontaneous activity, and their interaction at each participant. Using 46 young and cognitively normal participants performing an event-related perceptual speed task inside the fMRI scanner, we showed that there is no significant interaction between the spontaneous and the task-evoked activity either as negative BOLD response in the DMN or as positive BOLD response in the DAN. We replicate our findings in a separate dataset with 24 young and cognitively normal participants performing a different event-related task inside another fMRI scanner. Our results suggest that the presence or absence of coherent spontaneous activity in the DMN, or DAN has negligible effect on the magnitude of the BOLD response and vice versa, which consequently means that the fMRI signal is a linear superposition of spontaneous and task evoked activity.

## 1. Introduction

One of the long-standing yet unanswered questions in neuroscience is whether spontaneous brain activity interacts with task-evoked activity. Spontaneous brain activity represents the intrinsic or ongoing activity of the brain that optimally is measured during rest (Biswal et al., 2010; Fox et al., 2005; Greicius et al., 2003), while task-evoked brain activity reflects the specific neural responses stimulated by external stimulus or task (Cabeza & Nyberg, 2000). Interacting spontaneous and task-evoked activities means that the amplitude of the spontaneous activity at the time of external stimulation significantly alters the magnitude of the co-localized task-evoked brain response. The answer to such a seemingly simple question turns out to be extremely difficult mainly because the neural processes underlying spontaneous brain activity are continuously functioning at all times, even during task performance (Buckner et al., 2008; Gusnard & Raichle, 2001; Raichle, 2015), which makes their disentanglement from the co-localized task-evoked activity really a difficult task. Therefore, direct and effective measurements of spontaneous brain activity during task performance are challenging, if not impossible. Researchers often use proxy measurements, such as the magnitude of the signal during the pre-stimulus period (He, 2013; Huang et al., 2017), to address this issue. However, using the pre-stimulus interval as a proxy measurement for spontaneous activity is suboptimal because the spontaneous activity is an ongoing activity and continuously changing; thus, the magnitude of the spontaneous activity during the pre-stimulus period does not determine whether it is going to increase or decrease at the stimuli onset or afterward. That is why more recent studies have tried to use the signal’s phase to predict its direction (Huang et al., 2017).

Despite the difficulties, the answer to this question has been shown to have many implications in several aspects of the neuroscience field (Nierhaus et al., 2009; Northoff et al., 2010). For instance, it will help in 1) understanding the intrinsic versus extrinsic functional architectonics of the human brain’s large-scale networks (Cole et al., 2014; Mennes et al., 2013), 2) addressing the long-standing concern about the trial-by-trial averaging (brain response variability), a customary practice for detecting task-evoked activity in electrophysiological recording (Arieli et al., 1996), fMRI (Becker et al., 2011; Nierhaus et al., 2009; Sadaghiani et al., 2010), and EEG (Mayhew et al., 2013) studies, 3) understanding the lack of difference in total brain activity, blood flow, or glucose consumption between rest and task. 4) Understanding the role of spontaneous activity in inducing trial-by-trial variability in evoked response (Arazi et al., 2017; Fox et al., 2006; Ito et al., 2020; Wolff et al., 2019, 2021). Most existing studies investigating the interaction between spontaneous and task-evoked activities focus on answering one of these questions, influencing their method of choice for such an investigation. For example, the trial-by-trial variabilities in the task-evoked brain responses have been hypothesized to be the result of a strong co-existing and co-localized spontaneous brain activity, which is superimposed with the task-evoked activity to give the total neuronal activity without any interaction; thus, justifying the trial-by-trial averaging as a method for extracting task-evoked activity.

The advent of functional magnetic resonance imaging (fMRI) and the recent advancements in acquisition technology and post-processing techniques have revolutionized our understanding of human brain functionality, particularly its system-level functional architecture. Despite these breakthrough advances, like other functional imaging modalities, fMRI also lacks an effective and direct measurement of the brain’s spontaneous activities during task performance. Task-evoked activities are often measured by relying on the timing of the task and measuring time-locked signal changes due to each stimulus, whereas inter-regional correlations in fMRI signals, often referred to as functional connectivity, are usually used to measure spontaneous activity. While we can use data-driven and multivariate techniques such as spatial independent component analysis (ICA) to extract the time course of the spontaneous activities during task-based fMRI, the task-evoked co-activated/deactivated regions are often co-localized with main functional connectivity networks such as DMN and DAN which makes their extracted time course to be partially contaminated with task-evoked activities (Calhoun et al., 2001; Smith et al., 2012). In this work, the time course of the spontaneous brain activity of each network is extracted by orthogonalizing the task-evoked BOLD response and the network’s ICA time course. Therefore, for each participant we can use a voxel-wise multiple regression analysis to model the fMRI signal with the obtained time course of the spontaneous activity, predictor of task-evoked activity (convolved double-gamma HRF with task timing), and their interaction. The obtained voxel-wise regression coefficient for all participants will be fed into a second-level analysis to determine any interaction between spontaneous and task-evoked brain activity. Our results indicate that in both slow and fast-paced event-related fMRI design, there is no interaction between the task-evoked activity and spontaneous activity in neither of the DMN nor DAN regions.

Finally, we demonstrate theoretically, with simulation, and with actual fMRI data that orthogonalizing ICA time course and task-evoked response has no effect in the interaction term within the multiple regression analysis. Essentially, we will show that orthogonalizing or not orthogonalizing the time course of the two obtained brain activities have no effect on the interaction term. We first show this by reformulating the regression equations with the non-residualized variables and show the interaction term remains unchanged. Then, using simulation we show that orthogonalizing the variables does not change the regression coefficient for interaction. Third, we repeat the analysis without orthogonalization and show the interaction results remain statistically equivalent. Altogether, our results provide evidence that there is no significant interaction between spontaneous brain activity and task evoked activity.

## 2. Methods

### Recruitment and Participants

This study utilized two sets of fMRI data from two different cohorts scanned with different scanners and fMRI task paradigms at two separate institutes. Dataset I included 46 healthy young subjects (mean age ± SD = 28.32 ± 5.54, female/male =22/24, all right-handed), which were recruited using a random market mailing approach within a 10-mile radius of Weill Cornell Medical Center (WCMC). Dataset II had 24 healthy young subjects (mean age ± SD = 24.75±3.14, female/male = 17/7, all right-handed) recruited using a random market mailing approach within a 10-mile radius of the Columbia University Irving Medical Center. For both datasets, the subjects were compensated for their time spent on this project, all subjects gave their informed consent before the scanning sessions, and the experimental design and subject recruitment procedures were approved by the Institutional Review Board in the corresponding research center.

### Experimental Design for fMRI Tasks

The first dataset employed an event-related fMRI paradigm wherein subjects had to perform a perceptual speed task referred to as *pattern comparison* (Razlighi et al., 2017). In this task, two abstract figures consisting of a varying number of lines connecting at different angles are presented alongside one another. Figure 1a illustrates two examples of this task stimuli. Participants are instructed to indicate whether the figures are the same or different by using a differential button press using the right hand (index finger used for indicating identical figures). The maximum allowable response time was 4 seconds, and the onset of the stimuli was jittered by applying a randomized inter-stimulus interval (ISI) generated from a uniform continuous distribution in the range of 1.0 – 4.0 second. The events were randomized once, and the same sequence was used for all the subjects. Participants were administered one run consisting of 74 trials completed in a scanning time of ten minutes. The timing of this task throughout the whole 10-minutes scan time are shown in Figure 1b. They were instructed to respond to each stimulus accurately and as soon as possible. The task was projected to a screen located at the end of the scanner and participant were laying down supine inside the scanner and were seeing the screen using a mirror located on the head-coil. All participants were trained outside scanner to perform the task comfortably.

**Figure 1:**
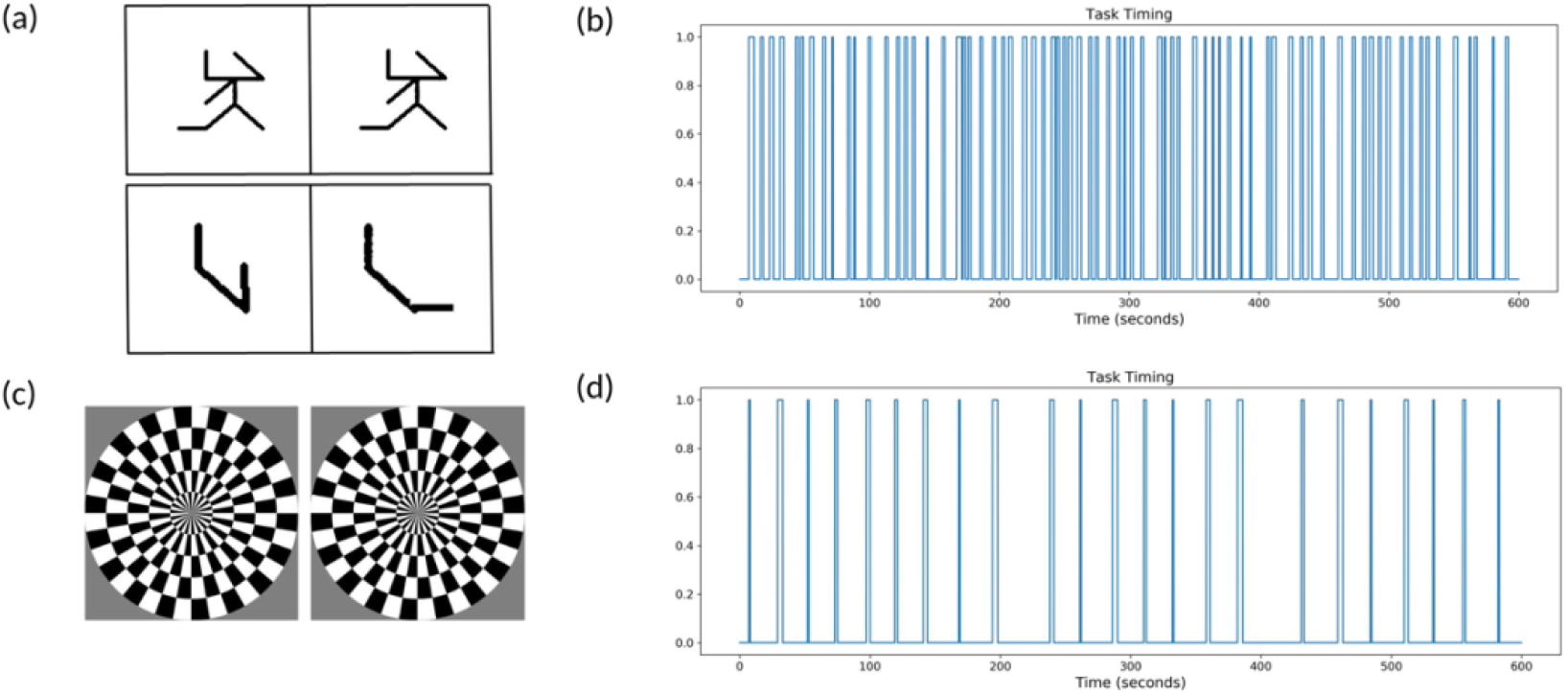
The event-related fMRI tasks paradigm used in the current study. (a) Two typical events of the pattern comparison task where participants had to decide whether they were the same or different (b) The timing of the 74 jittered events in the pattern comparison task used in Dataset I (c) The two alternating circular checkerboard visual stimuli alternating with 6Hz to strongly stimulate participants visual cortex (d) The timing of the 25 jittered events for circular checkerboard task in Dataset II

The second dataset had participants presented with a circular checkerboard flashing at 6 Hz and 100% contrast at the center of their field of view. This event-related fMRI paradigm had a longer ISI, which were sampled from a normal distribution ranging from 18 to 26 seconds. Event durations were selected randomly from a fixed set {0.5, 1, 2, 3, 4} seconds; but counter balanced to include five events of each duration adding up to 25 events presented in total. The events were randomized once, and the same sequence was used for all the subjects. The participants were asked to press a button twice with their right index finger at the end of each visual stimulus to ensure they were engaged in the task. Figure 1c shows the two circular checkerboard that they alternatively presented to the participants as the strong visual stimuli. The task was projected to a screen located at the end of the scanner and participant were laying down supine inside the scanner and were seeing the screen using a mirror located on the head-coil. All participants were trained outside scanner to perform the task comfortably. The timing of the task throughout 10-minutes of scan are illustrated in Figure 1d. As it seen the replication task has much longer ISI making this task relatively slower than the pattern comparison task.

### MRI acquisition parameters

Dataset I were acquired by a 3.0 Tesla Siemens Magnetom Prisma scanner and 64-channel head-coil using a T_2_^*^-weighted echo planar imaging (EPI) multiband sequence [TR/TE=1008/37 ms; flip angle=52°; FOV= 187.2 × 187.2 mm; matrix-size = 104×104; voxel-size = 2×2×2 mm^3^; 72 axial slices; multiband factor = 6]. The duration of fMRI scan was 10 minutes and 5 seconds (600 volumes). An accompanying T_1_-weighted magnetization-prepared rapid gradient-echo (MPRAGE) structural image was collected [TR/TE = 2400/3 ms; flip angle = 9°; FOV= 256 × 256 mm; matrix-size = 512 × 512; voxel-size = 0.5 × 0.5 × 0.5 mm; 320 axial slices] for localization and spatial normalization of the functional data in each participant.

Dataset II are acquired by another 3.0 Tesla Siemens Magnetom Prisma scanner and 32-channel head-coil using a T_2_^*^-weighted EPI multiband sequence [TR/TE=1000/30 ms; flip angle=62°; FOV= 200 × 200 mm; matrix-size = 100×100; voxel-size = 2×2×2 mm^3^; 64 axial slices; multiband factor = 4]. The duration of fMRI scan was 10 minutes (600 volumes). An accompanying T_1_-weighted MPRAGE structural image was collected [TR/TE = 2300/3 ms; flip angle = 8°; FOV= 256 × 256 mm; matrix-size = 256 × 256; voxel-size = 1 × 1 × 1 mm; 196 axial slices] for localization and spatial normalization of the functional data in each participant.

### Preprocessing and statistical analysis of fMRI data

All fMRI data were analyzed using FSL (V6.0.4) (Jenkinson et al., 2012) and in-house-developed packages. The preprocessing of the fMRI data involved the following steps. A slice-timing correction was applied to the raw fMRI time series to account for the difference in the acquisition delay between slices. At the same time, the motion parameters were estimated on raw fMRI scans using rigid-body registrations performed on all volumes having the first volume as the reference. Geometric distortion was corrected by *topup* technique as provided in FSL (Smith et al., 2004) using reference volume from another fMRI scan acquired with opposite phase encoding direction. Then, the estimated motion parameters were applied to the slice timing corrected fMRI data to get both temporally and spatially re-aligned fMRI time series. To enhance the precision in the localization of the statistically significant voxels, a spatial smoothing with small kernel size (FWHM = 3mm) was applied. A temporal high-pass filter (>0.01 Hz) was applied to the fMRI time series to account for the slow scanner drift in the data. Spatial normalization was performed by a rigid-body transformation of the first fMRI volume to its T_1_-weighted structural scan of the same participant and then by non-linear registration of the structural image to the MNI template.

The preprocessed fMRI data were then fed into the FSL multivariate exploratory linear optimized decomposition into independent components (MELODIC) to perform subject-wise ICA with the number of IC being determined automatically using Laplace approximation to the Bayesian evidence of model order. The ICs are visually inspected, and the two IC are identified which spatially correspond to the DMN and DAN regions. The operator that was identifying the ICs (first author) was blind to the participant’s demographics and health status. The IC with spatial pattern (Z>3) covering all three main nodes of the DMN (posterior cingulate, medial frontal gyrus, and bilateral angular gyrus) was selected as the DMN IC. Similarly, the IC with spatial pattern (Z>3) covering bilateral inferior parietal, middle temporal gyrus, and frontal eye fields was selected as the DAN. The timeseries of the identified ICs was orthogonalized with respect to the predictor of the task-evoked activity to obtain the predictor of spontaneous activity in the subject-level multiple regression analysis.

The subject-level voxel-wise multiple regression analysis was performed by modeling the each voxel’s fMRI timeseries with three predictors of interest; 1) The task-evoked activity which was obtained by convolving the canonical double-gamma HRF with the timing of the stimuli (zero-one boxcar series shown in Figure 1), 2) The spontaneous activity which was obtained by performing subject-wise ICA on the same subject’s fMRI data and orthogonalizing it in respect to task-evoked activity, and 3) Their interaction; as well as 6 motion parameters as nuisance. Equation 1 show the final model run for each voxel.

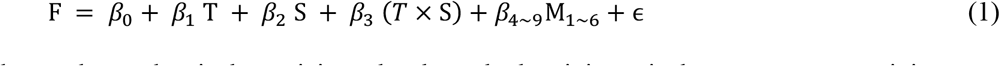

where *F* is the fMRI signal at each voxel, *T* is the anticipated task-evoked activity, *S* is the spontaneous activity, *(T × S)* is the interaction term modeling the modulating effect between two forms of activity, ϵ is the unexplained variance in the fMRI signal, and M_1_ to M_6_ are the six motion parameters obtained during spatial realignment. Running the voxel-wise multiple regression analysis given in equation 1 results in three different statistical parametric maps for *β*_*1*_ ∼ *β*_3_ for each participant. Group-level analysis is performed by feeding the obtained parametric maps into a second-level hierarchical regression analysis using FLAME, and a mixed-effects modeling technique implemented in the FSL. The results after cluster-wise multiple comparisons correction will determine the voxels activated during the stimulation, the voxels that show significant spontaneous activity for the network being investigated, and whether the spontaneous activity modulates the task-evoked response in that voxel.

### Effect of orthogonalizing variables on their interaction

In the previous section, we obtained the timeseries of spontaneous activity by orthogonalizing the identified network’s ICA timeseries in respect to the task-evoked activity. This might raise a question that what is the effect of such orthogonalization on the interaction term. In this experiment, we aim to show that in general orthogonalizing two variables in a multiple regression analysis (such as Equation 1) has no effect on the interaction term. Essentially, we are trying to show that our findings are not the consequence of orthogonalizing the two brain activities and even if we omit the orthogonalization step our findings will not change. First, we start by reformulating the Equation 1 with the original (non-orthogonalized) spontaneous activity timeseries Ŝ = *w*_0_ + *w*_*1*_*T* + *S*, where Ŝ is the original timeseries, *T* is the same anticipated task-evoked acticity, and *w*_0∼*1*_are the regression coefficient for the orthogonalizing linear model. Now, replacing *S* in Equation 1 with Ŝ – *w*_0_ – *w*_*1*_*T*, and simplifying the equation will give the following equations.

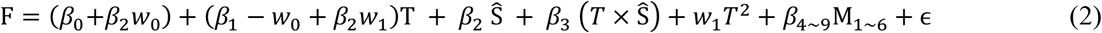

As it seen, the interaction term still has the same coefficient, *β*_3_, as it had in the Equation 1, even though other coefficients may have changed. This is evident that orthogonalizing the two variables in the multiple regression analysis has no effect on the interaction term.

Next, we perform a simulation test to demonstrate that orthogonalizing the independent variables in a multiple regression analysis has no effect on the interaction term. We first show this effect in interacting variables and also on a pair of non-interacting variables to make sure that orthogonalizing two variables does not induce false-positive interaction. To this end, we first simulated two random variables from a multivariate normal distribution with zero mean and unit variance which are also correlated with *ρ* = 0.25. We realized 600 sample points from each variable to match the length of the timeseries in our empirical data. Using the two correlated variables and equation 1, we then synthesized two sets of 100000 random timeseries by adding 50% white Gaussian random noise to each timeseries. In the first set of simulation, we made the two variables interacting by setting *β*_*1*_ = *β*_*2*_ = 1, & *β*_3_ = 1 whereas in the second set of the simulation they were not interacting *β*_*1*_ = *β*_*2*_ = 1, & *β*_3_ = 0. The result of these simulation were two sets of 100000 timeseries. In the first set, the effect of each variable on the synthesized timeseries were dependent on the value of the second variable whereas in the second set the two variables were not interacting even though the actual variables were correlated. Note that the correlated variables do not necessarily have an interaction effect on a dependent variable. Next, for each set of the 100000 synthesized timeseries, we use both original and orthogonalized variables to model the synthesized timeseries and obtained all the regression coefficients. For both sets, we showed that the estimated regression coefficient for the interaction term using the original variables 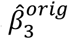, is statistically equivalent to the estimated coefficients for interaction term of the orthogonalized variables 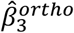. This is done by comparing the mean of the two coefficients using two sample t-test.

Finally, we reanalyze our fMRI data for both datasets without orthogonalizing the spontaneous and task evoked timeseries and show that the obtained statistical map for interaction is statistically the same as the one obtained previously. We perform the same analysis as it is explained above, only with the difference that the spontaneous activity was not orthogonalized with respect to task-evoked activity. The obtained statistical parametric maps for the interaction term then compared with the one we obtained before and any statistical differences between two maps are assessed using a voxel-wise two-sample student t-test.

### 3. Results

Table I shows the participant demographics in both datasets. Our samples have no significant gender differences for the first cohort (*χ*^*2*^=0.087, p > 0.77), but there is significant gender difference in the replication dataset (*χ*^*2*^=4.17, p < 0.041). Both cohorts are young and healthy individuals, but the replication cohort in average is 4.75 years younger that the first cohort. Each participant in both cohorts has been trained about the task, and they all learn to do perform the task outside the scanner. The task performances (accuracy, response time and the time-out rate) during the scanning are listed in Table I. As it seen, both cohorts learn the task very well as they perform the task with very high accuracy and low time-outs rate. Even though both tasks are considered easy, the performance on the replication task (Circular Checkerboard) is naturally better than the pattern comparison, as it is still easier than the pattern comparison task.

**Table 1:** Demographics and task performance of the two datasets.

|  | Number of Subjects | Age (years) | Sex (M/F) | Accuracy (%) | Response time (second) | Time outs (%) |
| --- | --- | --- | --- | --- | --- | --- |
| Dataset I (WCMC) | 46 | $28.32 \pm 5.54$ | 24/22 | $97.0 \pm 3.65$ | $1.823 \pm 0.305$ | 1.32 |
| Dataset II (CUIMC) | 24 | $24.75 \pm 3.14$ | 17/7 | $99.0 \pm 2.38$ | $0.645 \pm 0.088$ | 0.33 |

The design and timing of the stimuli in our fMRI experiment for the perceptual speed task used in Dataset I and the flashing circular checkerboard task in the replication study are shown in Figure 1. As it seen, the fMRI task paradigm in the first dataset is much faster than the task paradigm in dataset II. We start by presenting the result for the first Dataset and then show the results from the replication dataset.

### Interaction of Task-evoked and Spontaneous Activities

The aim of this experiment is to determine whether there is an interaction between task-evoked and spontaneous brain activity. In each participant, ICA is used to extract the time course of the two well-recognized brain networks (DMN and DAN) activities while the participant was performing a simple perceptual speed task (pattern comparison). The obtain timeseries are orthogonalized with respect to the anticipated task-evoked BOLD response to get a unique measurement of the brain spontaneous activity for these two networks. In each participant, a multiple linear regression is used to model each voxel’s fMRI timeseries with the anticipated BOLD response, the obtained unique measurement of spontaneous activity, and their interaction. As given in equation 1, the six motion parameters are also added as the nuisance variables to the model. The statistical parametric maps (regression coefficients) of each of the three variables of interest are fed to the group levels analysis. Figure 2 shows the results of the group level analysis for both DMN and DAN after performing cluster-wise multiple comparison corrections with t-statistics threshold of 4 and p-value of 0.01 for all obtained patterns of brain activity. The pattern of task-evoked positive BOLD response (activation) and negative BOLD response (deactivation) in response to pattern comparison tasks are given in Figure 2a and 2d. As it is seen in Figure 2, the voxel-wise t-statistical significance of the group-level activations are overlaid on a semi-inflated cortical surface of the brain with hot colors (red to yellow) and deactivations with cold colors (dark blue to light blue). As it is expected, these two patterns are essentially the same, as they are just depicting the regions that are responsive to the task. As it is seen, the pattern comparison task generates a positive BOLD response (activation) in the visual cortex, left hemisphere sensory motor (since all participants respond with right hand) and regions that overlap with DAN and a negative BOLD response (deactivation) in the DMN regions. In addition, we observe significant activations in the fronto-parietal regions and significant negative BOLD response in bilateral ventricular regions. The pattern of spontaneous brain activity is also depicted using the t-statistical significance of the group-level analysis for spontaneous activity and overlaid on a semi-inflated cortical surface in figures 2b and 2e for DMN and DAN respectively. It is noteworthy that even after orthogonalizing the DMN and DAN timeseries in respect to the task-evoked BOLD response, the regions of the DMN and DAN are demonstrating very strong co-activities that are un-related to the task-evoked activities. The magnitude and significance of co-activations due to our orthogonalized measurement of spontaneous activity is almost twice as the one observed for task-evoked activity. As expected, the co-activation pattern for the DMN is corresponding to the know DMN regions and the co-activation pattern for the DAN corresponding to the know DAN. In fact, the patterns of activity for both task-evoked and spontaneous response are very similar to the patterns obtained by modeling the fMRI timeseries separately and univariately with each activity. Figure S1 illustrates the results of three separate analysis each modeling fMRI timeseries with pattern of BOLD response (Figure S1a) and the spatial pattern of the DMN co-activation (Figure S1b), and the spatial pattern of the DAN co-activation (Figure S1c).

**Figure 2:**
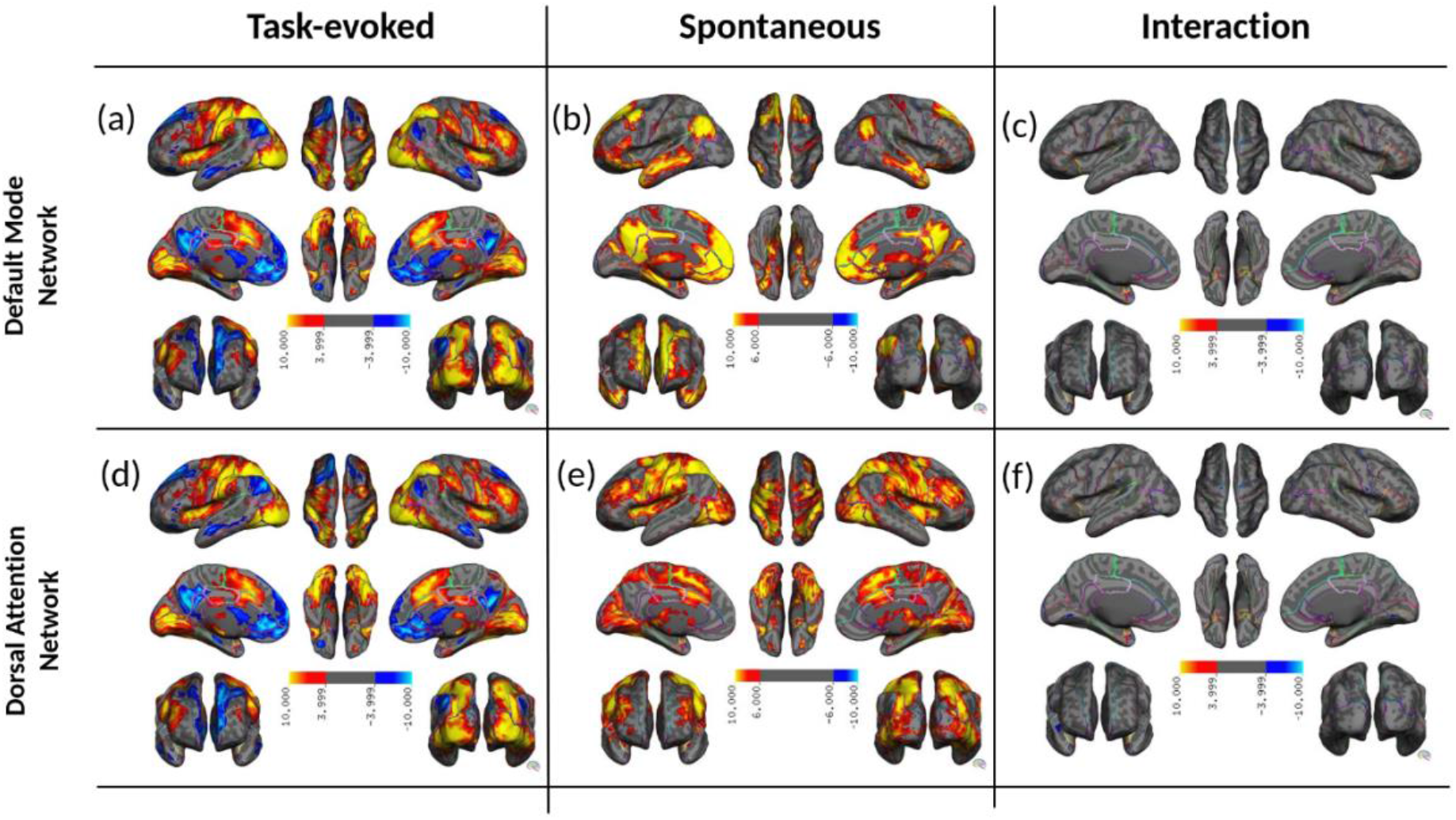
Spatial patterns of co-activated/co-deactivated voxels due to a&d) task-evoked activities, b) spontaneous activity of the DMN, e) spontaneous activity of DAN, and c&f) their interaction for pattern comparison task in dataset I. The significantly activated/deactivated voxels are color-coded with heatmap and overlayed on a semi-inflated surface of the cortex. The volumetric version of the figures depicted in Figure S5 for DMN and in Figure S6 for the DAN.

Finally, the spatial pattern of the voxels showing a significant interaction between their task-evoked and spontaneous activities are depicted in Figures 2c and 2f for DMN and DAN, respectfully. These results suggest that at least for DMN and DAN there is no interaction between task-evoked activity and spontaneous activity. This is interesting because we identified a significant spatial overlap between task-evoked and spontaneous activity in DMN as well as in DAN. Next, we replicate our finding using a different cohort of participants scanned on different scanner and using different fMRI task paradigm.

### Replication with Different cohort and fMRI task Paradigm

The aim of this experiment is to replicate our finding in the previous section using a slower fMRI task paradigm to rule out the possibility that the absent of interaction might be due to fast nature of our fMRI design in pattern comparison task. Our replication study has a task paradigm which is much slower in the rate of events stimulation (25 events versus 74 events in 10 minutes of scanning) and has a completely different cohort which is scanned in different institutes. The same pre-processing pipeline is used to process the fMRI data in the replication study and the same statistical model as given in equation 1 is used to model the voxel-wise timeseries. Figure 3 shows the results of the group-level analysis after cluster-wise multiple comparisons correction with a t-statistics threshold of 3 and p-value of 0.01 for all obtained patterns of brain activity. Figures 3a and 3d depict the spatial pattern of task-evoked activity by overlaying the voxel-wise t-statistics on a semi-inflated cortical surface of the brain. Activities in visual cortex and primary motor area in the left hemisphere are evident, however there are also activation in the area of the DAN. On the other hand, deactivations on the DMN are also seen in these figures, even though they are much weaker than the one observed in the first experiment. This might be because this task has only 25 events which is almost one-third of the number of events in the first experiments, and almost half of the number of participants (24 versus 46 in the first experiment). The activations on visual cortex are not weaker, because the task presented here are primarily a visual stimulus (flashing checkerboard) which is specifically designed to induce maximal visual response. Figure 3b and 3e depicts the spatial pattern of spontaneous activities for DMN and DAN, respectively. As it is seen, both DMN and DAN timeseries generated activities in the same regions often reported for these two networks in the literature. The spatial patterns obtained for DMN and DAN are similar to the ones we had obtained from the first dataset although they appear to be weaker in strength which make sense due to aforementioned reasons. Figure S2 illustrates the pattern of both task-evoked activity (Figure S2a) and spontaneous activities (Figure S2b for DMN and Figure S2c for DAN) when they are obtained using three separate univariate analyses with a single variable of interest at a time. As seen, the activity pattern is very similar whether we include all three variables of interest or one variable at a time. Finally, the interactions are depicted in Figures 3c and 3f, where we found no interaction between the two activities for both DMN and DAN, thus replicating our results obtained from the first dataset.

**Figure 3:**
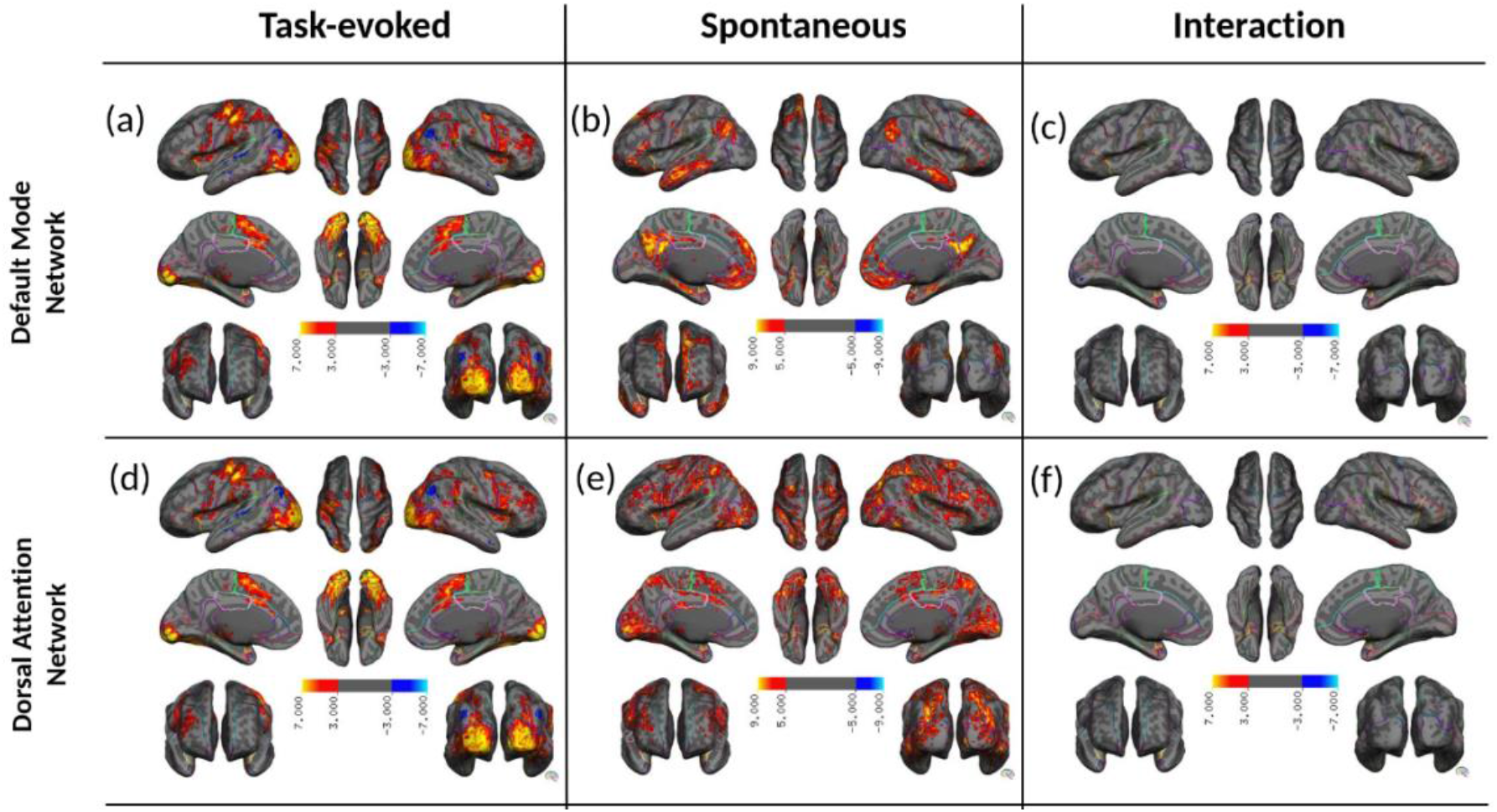
Spatial patterns of co-activated/co-deactivated voxels due to a&d) task-evoked activities, b) spontaneous activity of the DMN, e) spontaneous activity of DAN, and c &f) their interaction for circular checkerboard task in dataset II. The significantly activated/deactivated voxels are color-coded with heatmap and overlayed on a semi-inflated surface of the cortex. The volumetric version of the figures depicted in Figure S7 for DMN and in Figure S8 for the DAN.

The group level statistics for the first cohort are shown with different ranges as compared to the second cohort due to differences in image acquisition, number of subjects as well as the change in the number of events presented in the task for the two cohorts. We also note that the color bar is not consistent for spontaneous activity as compared to the task-evoked activity. This can be due to two reasons: (i) task-evoked activity is present only for short durations over a ten minute scan while spontaneous activity spans the full ten minute period hence leading to stronger signals in Figure 2b and Figure 2c as compared to Figure 2a and 2d, (ii) spontaneous activity is present with or without the task and hence one could hypothesize its presence being fundamental to the brain’s core processes; thus it has stronger signal strength. Next, we aim to address the issue associated with orthogonalizing the task-evoked and spontaneous activities regressors and show orthogonalizing the two variables, in general, has no effect on their interaction term.

### Orthogonalizing two variables does not alter their interaction

In this experiment we aim to show that in the model shown in Equation 1, orthogonalizing the two variables does not significantly affect the interaction terms. We have already shown this by reformulating equation 1 in terms of non-orthogonalized variables and derived equation 2 where the interaction term’s coefficient remained intact after orthogonalizing the two variables. Here we show the results of the same finding using simulation and re-analyzing the fMRI data with variables not being orthogonalized.

As described in the method section, using the two correlated gaussian random variables (*ρ* = 0.25) and equation 1, we synthesized two sets of 100000 timeseries and added 50% white Gaussian random noise. In the first set of simulations, we made the two variables interact by setting *β*_*1*_ = *β*_*2*_ = 1, & *β*_3_ = 1, whereas in the second set of the simulation, they were not interacting *β*_*1*_ = *β*_*2*_ = 1, & *β*_3_ = 0. Next, for each set of the synthesized timeseries, we used original and orthogonalized variables to model the synthesized timeseries and obtained all the regression coefficients. For the first set, we found that the estimated regression coefficient for interaction using the original variables 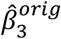, is statistically equivalent to the estimated coefficients for the interaction term of the orthogonalized variables 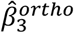, by rejecting the null hypothesis: 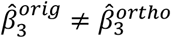(t=0.165, p>0.87), and similarly for the second set (non-interacting), by rejecting the null hypothesis: 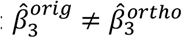, (t=0.51, p>0.61). Altogether, these simulations show that, whether we have interacting or non-interacting variables, orthogonalizing the two variables generates the same coefficients for the interaction terms in a multiple regression analysis.

Finally, we have reanalyzed both datasets used in this work without orthogonalizing the DMN and DAN ICA timeseries with respect to task-evoked activities. Note that we do not claim that the ICA timeseries obtained from task-based fMRI scans are an accurate measurement of brain spontaneous activity, instead we aim to show that orthogonalizing the ICA timeseries does not have any effect on our obtained results and even without orthogonalizing them, there will be no interaction between the two brain activities. This experiment is mainly performed to address the concern that our finding might be the results of orthogonalizing two predictors of brain activities. Figure S3 shows the results of this analysis for database I. As it seen, Figures S3a and S3d show the mean of the regression coefficients for interaction term (*β*_3_) for every voxel in the brain overlaid on a semi-inflated surface of the cortex when investigating DMN and DAN, respectfully. The values are color-coded with heatmap. Note that the statistical significance of these regression coefficients is essentially depicted in Figures 2c and 2d after correcting for multiple comparisons correction. Figures S3b and S3e shows the same mean of the regression coefficients for interaction term but this time without orthogonalizing the two brain activity measurements. Finally, Figures S1c and S1f shows the statistical difference between the two regression coefficients at every voxel overlaid on a semi-inflated brain surface. The t-statistics are color-coded with heatmap. As it seen, we found no significant differences between the two sets of regression coefficients. This finding providing further evidence that orthogonalizing the two predictors of brain activities has no effect on the interaction term. Figures S4 provides the results of the same analysis performed on dataset II. Again, as it seen in figure S4c and S4f, there is no significant differences between the interaction regression coefficients obtained with and without orthogonalizing the two brain activity measurements. Altogether, these findings alleviate the concern about our results being the consequences of orthogonalizing variables.

## 4. Discussion

In this work we have used, for the first time, the time series of the functional connectivity networks obtained by subject-wise spatial ICA as the continuous measurements of that network’s spontaneous activity. Due to spatial overlap between functional connectivity networks and task-evoked co-activated regions, the obtained network’s timeseries from the task-based fMRI scans using spatial ICA were often contaminated with task-evoked variabilities (Calhoun et al., 2001; Smith et al., 2012). Thus, to have a unique measurement of the spontaneous activity, we orthogonalized the identified network timeseries with respect to the anticipated task-evoked activity. The orthogonalized network timeseries generated a spatial activation pattern that matched that network’s spatial pattern. We showed no significant interaction between the two activities using the orthogonalized time series for spontaneous activity, anticipated BOLD response for task-evoked activity, and their interaction in a voxel-wise multiple regression analysis. We analyzed two well-recognized functional connectivity networks (DMN and DAN). We replicated our findings using a much slower task paradigm (ISI > 18 sec compared to ISI < 4 sec for the first dataset). Finally, we have shown that orthogonalizing the predictors of the spontaneous and task-evoked activities in the multiple regression analysis does not affect the interaction term. We have demonstrated this by reformulating equation 1, then by simulation, and finally by re-analyzing both datasets using the non-orthogonalized time series.

A few studies have looked at the association between task-evoked and spontaneous activity focusing on the role of spontaneous activity on sensory perception. One study reported that the degree to which DMN was suppressed during stimulus epochs correlated positively with task-evoked activity in auditory and visual cortices (Greicius et al. 2004), suggesting that attention to stimuli or lack thereof can be deduced from passive tasks. Another study (Boly et al., 2007) reported a negative relationship between conscious perception of pain stimulation and baseline activity in the regions encompassing the DMN, suggesting that spontaneous activity modifies the conscious perception of the external world. Another study reported that the degree of task-evoked activity decline could be related to the variability of spontaneous activity, which significantly correlated with task performance accuracy, laying the groundwork for the behavioral significance of task modulation of functional connectivity variability (Elton & Gao, 2015).

It is important to note that the resting-state functional connectivity networks as an indicator of brain spontaneous activity have also been observed in the presence of tasks (Hasson et al., 2009), and they don’t seem to differ much from functional connectivity generated during tasks (Bolt et al., 2017). Several studies investigated the effect of task performance on large-scale brain functional connectivity networks, mainly seeking a task-induced reconfiguration in the brain’s functional architecture. A seminal work by Cole et al. (Cole et al., 2014) showed that the brain’s functional network architecture changes marginally during different cognitive tasks (close to 10%), which questions whether there is any change at across tasks. Using the Psychophysiological Interaction (PPI) approach, Di and colleagues showed that task-evoked activity modulated spontaneous activity in four different cognitive tasks in regions that were activated, deactivated, or had no activation to the task at all, leading authors to conclude that no clear relationships exist between task-modulated connectivity and regional task activations (Di & Biswal, 2019). Using activity flow models, Cole et al. showed that estimating task-evoked activity flow (the spread of activation amplitudes) from spontaneous activity allowed the prediction of task-evoked activations in a large-scale neural network model, thereby explaining the relevance of resting-state functional connectivity networks to cognitive task activations (Cole et al., 2016). A study done by Gratton and colleagues (Gratton et al., 2016) demonstrated that task-related activation and correlation-based functional connectivity measures, albeit being related, might index separable aspects of brain function. These results highlight the role of spontaneous activity on task-evoked processes, giving rise to thoughts that spontaneous and task-evoked activity might be intricately related. In this current report, we investigate whether spontaneous activity interacts with task-evoked responses in regions that are positively (activations) or negatively (deactivations) affected by the task, going beyond mere association between spontaneous activity and task-evoked activity and show that such thoughts of interactions are not valid.

Several studies have directly attempted to examine the interaction between task-evoked and spontaneous brain activity using trial by trial examination of fMRI signal as it has been reviewed in few articles (Nierhaus et al., 2009; Northoff et al., 2010; Sadaghiani et al., 2010). There are numerous challenges in performing these studies. For instance, 1) it is impossible to directly dissociate spontaneous and evoked activity from the recorded signal, as the spontaneous activity may continue to vary after stimulus onset, 2) it is difficult to disentangle whether up-coming task trial effecting spontaneous activity versus spontaneous brain activity effecting evoked response, 3) our knowledge about the spontaneous activity, its role in brain function, and its underlying neural processes are very limited, 4) the measurement of spontaneous activity, if not done carefully, can lead to spurious findings as reported previously (Logothetis et al., 2009). Despite these challenges, researchers used different approach to study the relationship between evoked and spontaneous brain activity. In general, the exiting approaches can be categorized into two main groups. The first group assumes that the underlying spontaneous activity can fully explain the observed trial-by-trial variability in evoked response. Therefore, the recorded response is a linear superposition of the task-evoked and spontaneous activities. Numerous studies used this approach to demonstrate that superposition holds between the two brain activities (Arieli et al., 1996; Azouz & Gray, 1999; Becker et al., 2011; Fox et al., 2006; Saka et al., 2010). The second group considered the magnitude of the recorded signal during the pre-stimuli period as the measurement of spontaneous activities (He, 2013; Huang et al., 2017; Mayhew et al., 2013). This group often found that the level of spontaneous activity measured during the pre-stimulus period is negatively associated with the magnitude of the evoked response suggesting that there is an interaction between spontaneous and evoked activity; thus, invalidating the superposition hypothesis (He, 2013; Huang et al., 2017; Mayhew et al., 2013). More strikingly, the recent report by Haung et. al, analyzed their data using the method proposed in the first group and found that there is no interaction between the two brain activities (Huang et al., 2017). Altogether, these studies highlight the shortcoming of both methods used for investigating the relationship between evoked and spontaneous activity as it has been reported previously (Logothetis et al., 2009). The method proposed here is a different way of looking at the interaction between task-evoked and spontaneous activity. Unlike the existing methods that look at trial-by-trial relationships, we assume that spontaneous brain activity is an underlying and continuous process, and it can be detected using a data-driven approach (i.e., ICA) to extract a covariance pattern from a multidimensional fMRI data. We show that for task-positive activation in the regions associated with the DAN or task-negative deactivation in the regions associated with the DMN, the impact of spontaneous activity on the task-evoked activity or vice vera is negligible.

### Correlation, Interaction, and Superposition

While the theoretical definitions for correlation, interaction, and superposition are clearly separate from each other, these terms are often used interchangeably in the literature which we believe is being a massive source of confusion in the field. For instance, correlated variables are not necessarily interacting, and interacting variables are not necessarily correlated. Superposition theorem states that in any linear system where more than one source is present, the response of the system is the sum of the responses obtained from each source considered separately. Here our sources are the task-evoked and spontaneous activities, and the responses are the MR signal changes due to those neural activities. The system is a set of complex processes that receive neural activations as the inputs and responds as changes in the brain hemodynamics and consequently fMRI signal. Essentially, the two sources can depend on each other, but the overall system they are inserted to can still be a linear system where superposition holds. In other words, the relationship between the two input signals does not determine the system’s linearity or the superposition’s validity. Interaction, on the other hand, indicates that if the relationship between task-evoked activity (T) and the fMRI signal (F) is changing based on the presence or absence of spontaneous activity (S), then S and T are interacting, and the superposition of S and T will not result in F, as it is shown clearly in equation (1). The main confusion comes when S and T become correlated, as shown in some previous studies (Becker et al., 2011; He, 2013; Huang et al., 2017), and often mistakenly interpreted as interacting.

This study aimed to show that there is no interaction between task-evoked and spontaneous activities; thus, the fMRI signal is a linear summation of the two responses (task-evoked and spontaneous activities) as it has been shown previously (Arieli et al., 1996; Azouz & Gray, 1999; Becker et al., 2011; Fox et al., 2006; Saka et al., 2010). We did not assess the association between task-evoked and spontaneous activity; therefore, our results should not be considered contradictory to the reported association between the two brain activities (Becker et al., 2011; He, 2013; Huang et al., 2017; Saka et al., 2010). Our analysis showed that the time course of spontaneous activities for DMN and DAN was correlated with the time course of the anticipated BOLD response. However, we feel this correlation is because spatial ICA cannot separate spatially overlapping processes as reported for these two networks (Parker & Razlighi, 2019; Smith et al., 2009). In addition, animal studies suggest that the brain’s spontaneous activities become coherent with the task-evoked response as long as the brain can anticipate the onset of external tasks (Cardoso et al., 2012; Sirotin et al., 2012). These reports may explain why, in a sparse but highly predictable task like the one used in Huang et al. 2017 (52<ITI <60), the spontaneous activity measured 10 seconds before the stimulus onset was shown to be associated with the task-evoked response. Both datasets we investigated were jittered with (1<ITI<4 sec) and (18<ITI<26 sec), which made it challenging for the participants to anticipate the upcoming stimuli accurately.

### Limitations and future works

In this study, we focused on two of the most studied large-scale brain networks, which have been shown to be spatially overlapping with task-based fMRI co-activated (DAN) and co-deactivated (DMN) regions (Kraus et al., 2021). However, many other brain networks remain to be investigated. In fact, Fox et al. (Fox et al., 2006) reported that super-position holds using the somatomotor networks, and surprisingly the same regions showed only a marginally significant interaction in Huang et al. study (Huang et al., 2017). These highlight the possibility that the interaction between task-evoked and spontaneous activity can differ across different brain networks or tasks. Therefore, future studies with and without sensory stimuli as well as cognitive tasks, are required to rule out these possibilities completely.

While our results demonstrate that there is no interaction between task-evoked and spontaneous activities, the time course of the extracted spontaneous activity for DMN and DAN was significantly correlated with the time course of the anticipated task-evoked activity for all the subjects in the first cohort (DMN: r *∼* −0.34 ± 0.13, and DAN: r *∼* 0.46 ± 0.16). The second cohort had similar results but less significant (DMN: r *∼* −0.15 ± 0.11, and DAN: r *∼* 0.098 ± 0.17. This correlation could be interpreted as the interaction between the two brain activities. There are many hypotheses explaining why evoked and spontaneous activities could be correlated (Becker et al., 2011; Fox et al., 2006; He, 2013; Huang et al., 2017; Saka et al., 2010). However, we feel that the observed correlation in our study is mainly due to the spatial overlapping evoked and spontaneous neural processes in these two networks, as it has been reported before (Smith et al., 2009). To further elaborate on this possibility, one could obtain the ICA coefficients for these two networks (DAN and DMN) from the resting-state fMRI scan of the same participant and use it to extract the network time series from the task-based fMRI. Another possibility is to use temporal ICA to extract the time series of the network, which is not sensitive to the spatially overlapping processes (Calhoun et al., 2001; Smith et al., 2012).

The results provided here give us a fresh perspective to look at spontaneous and task-evoked activity in the fMRI signal. We show that there is no interaction regardless of whether we control for the correlation between task-evoked and spontaneous activity time series or not. This suggests that the shared variance between task-evoked and spontaneous activity doesn’t play a role in their interaction. Unfortunately, we could not reanalyze our data using trial-by-trial variability approach due to the short ISI in our dataset (Huang et al., 2017). However, future studies with slower rate but fully jittered design are warranted to compare the results of our method with the one obtained using the trial-by-trial averaging and pre-stimulus period as the measurements of spontaneous activity.

## 5. Conclusion

The results presented in the current study provide evidence that there is no significant interaction between spontaneous and task-evoked activity, a fundamental scientific question in the field. We explained and demonstrated through simulation and re-analyzing the fMRI data that the two brain activities could be associated with each other, but that does not necessarily result in interaction between them. While we could not assess the validity of association between the two brain activities, we could show that independent of the presence or absence of any association between the two brain activities, there is no interaction between them, thus concluding that superposition holds between the two activities. Our finding provides further evidence that the fMRI signal is a linear summation of the MR signal changes due to spontaneous and task-evoked activity.

## Supplementary Material

**Figure S1:**
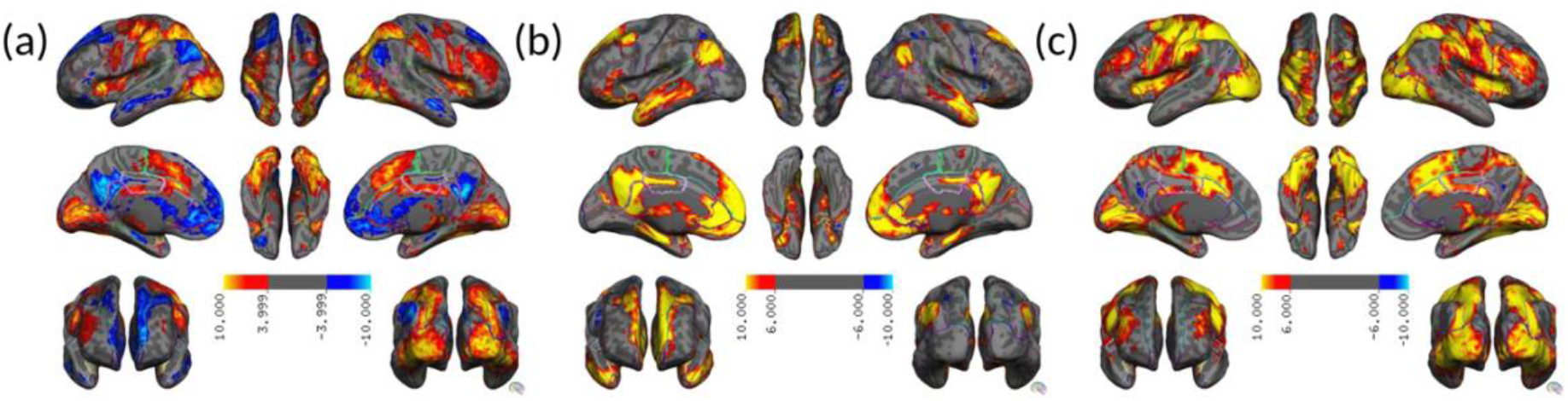
Results of reanalyzed the fMRI data in dataset I and pattern comparison task using three different univariate modeling; a) only with task-evoked predictors, b) only with obtained spontaneous activity predictor for DMN, and c) only with obtained spontaneous activity predictor for DMN. The significantly activated/deactivated voxels are color-coded with heatmap and overlayed on a semi-inflated surface of the cortex.

**Figure S2:**
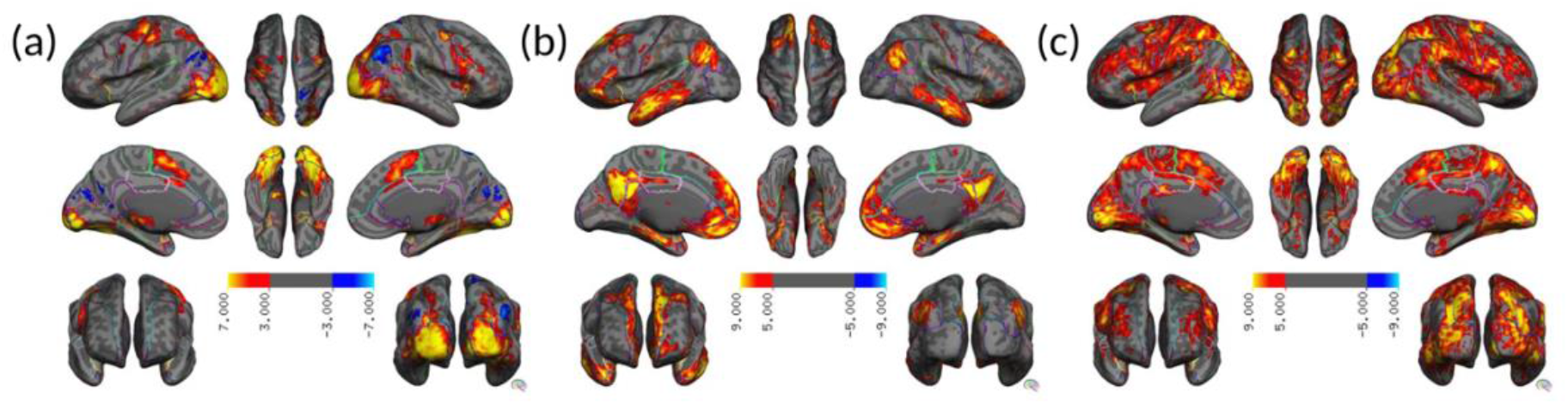
Results of reanalyzed the fMRI data in dataset II and circular checkerboard task using three different univariate modeling; a) only with task-evoked predictors, b) only with obtained spontaneous activity predictor for DMN, and c) only with obtained spontaneous activity predictor for DMN. The significantly activated/deactivated voxels are color-coded with heatmap and overlayed on a semi-inflated surface of the cortex.

**Figure S3:**
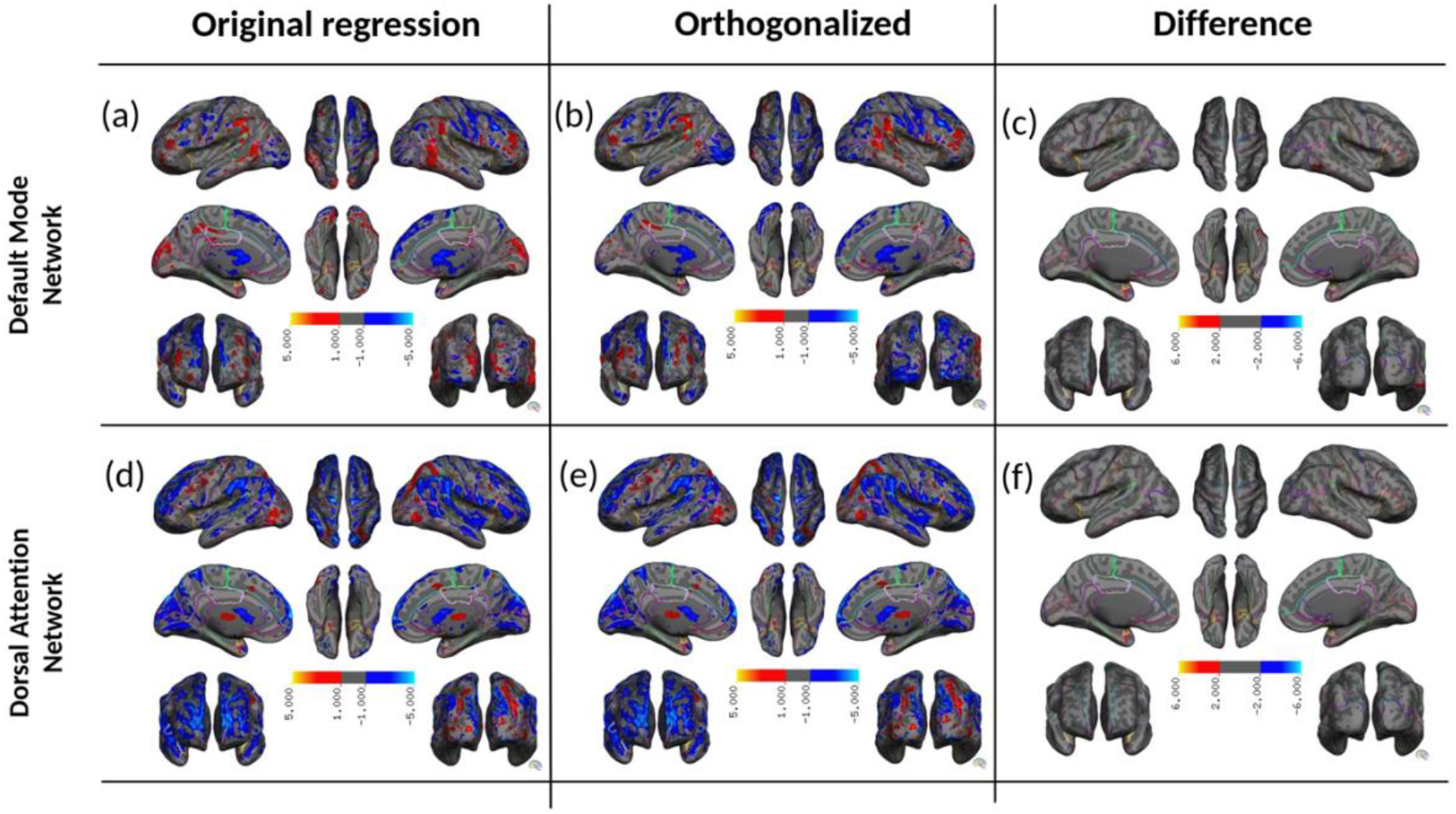
The spatial pattern of the mean regression coefficient for interaction term are color-coded with heatmap and overlayed on the surface of the cortex a&b) for DMN and d&e) for DAN for pattern comparison task in dataset I. a&d) depicts the voxel-wise interaction coefficient when original (non-orthogonalized) timeseries were used. b&e) depicts the voxel-wise interaction coefficient when orthogonalized timeseries were used. The significance of the voxel-wise difference in the mean of the two regression coefficients are obtain using student t-test and overlaid on the semi-inflated cortex c) for DMN and f) for DAN.

**Figure S4:**
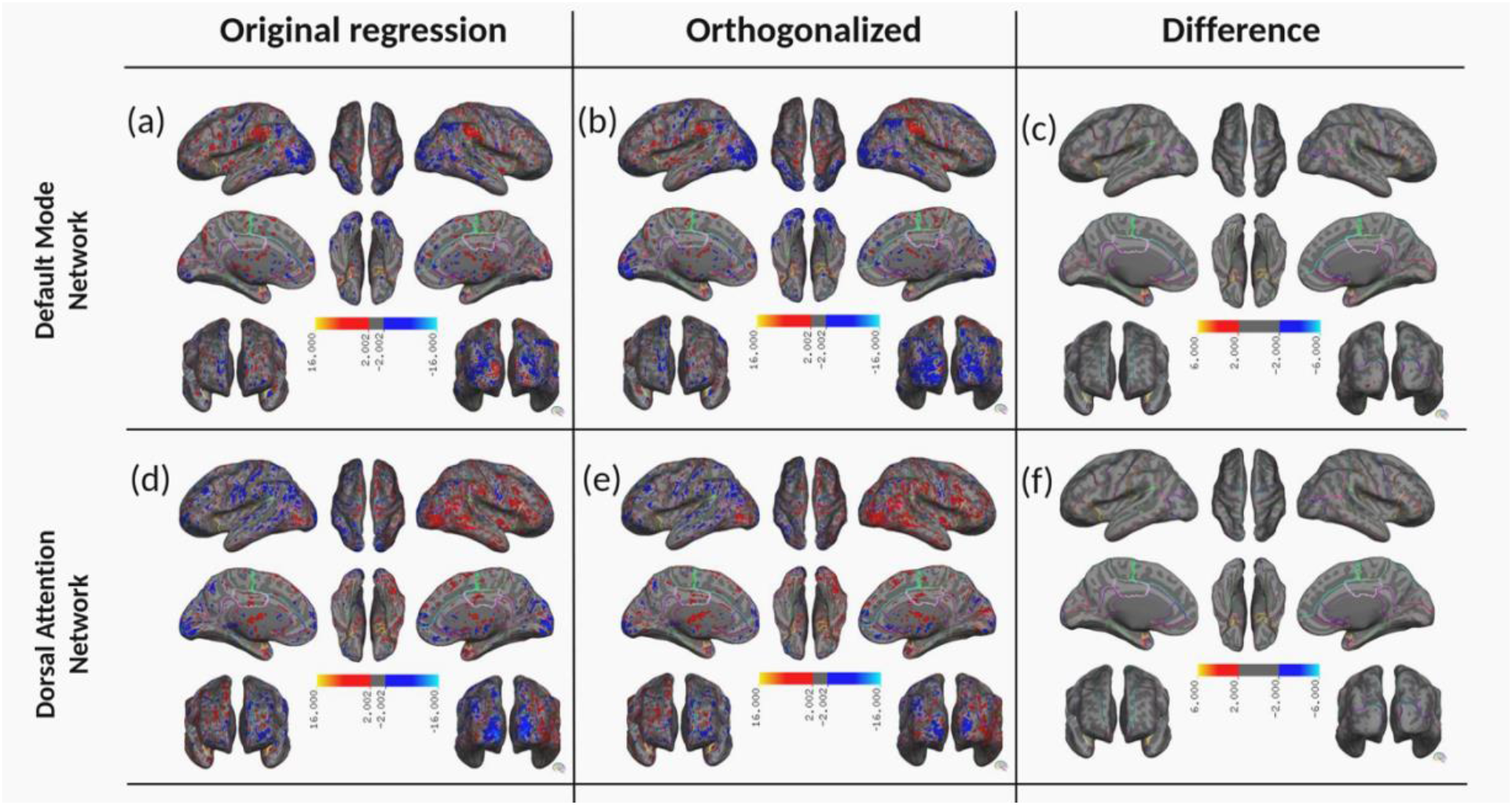
The spatial pattern of the mean regression coefficient for interaction term are color-coded with heatmap and overlayed on the surface of the cortex a&b) for DMN and d&e) for DAN for circular checkerboard task in dataset II. a&d) depicts the voxel-wise interaction coefficient when original (non-orthogonalized) timeseries were used. b&e) depicts the voxel-wise interaction coefficient when orthogonalized timeseries were used. The significance of the voxel-wise difference in the mean of the two regression coefficients are obtain using student t-test and overlaid on the semi-inflated cortex c) for DMN and f) for DAN.

**S5:**
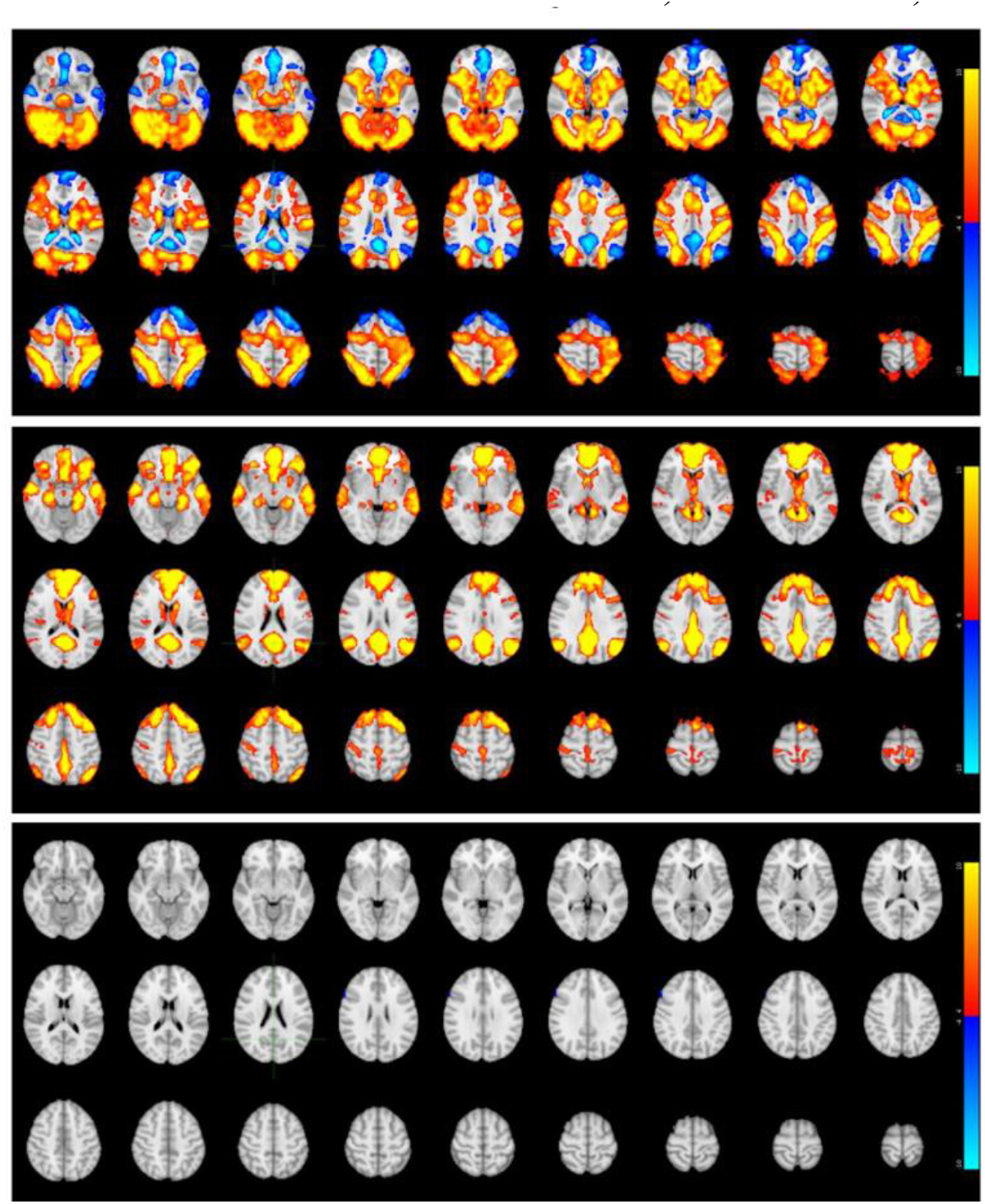
The volumetric version of the activation/deactivation patterns shown in Figures 1a, 1b, and 1c.

**S6:**
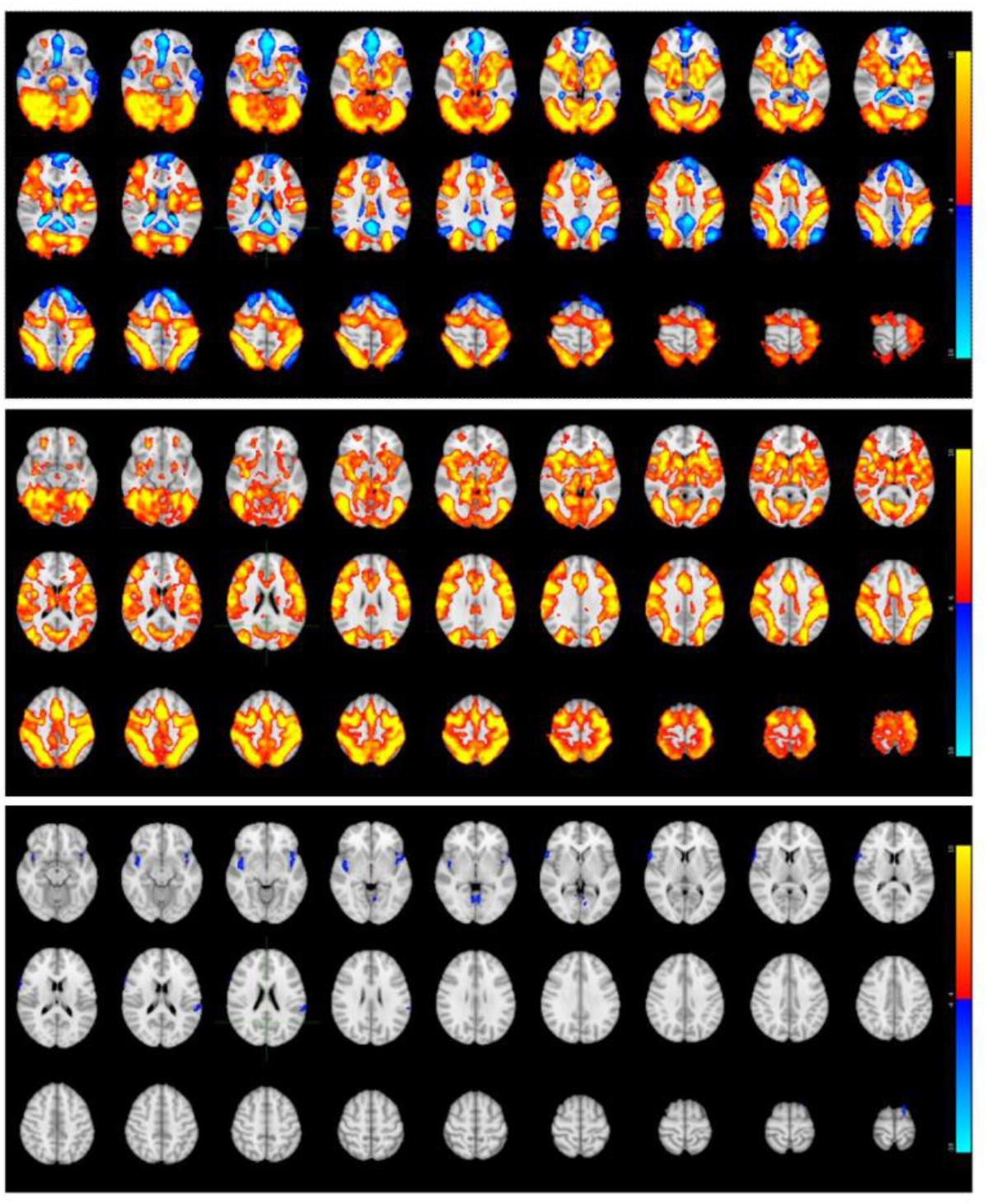
The volumetric version of the activation/deactivation patterns shown in Figures 1d, 1e, and 1f.

**S7:**
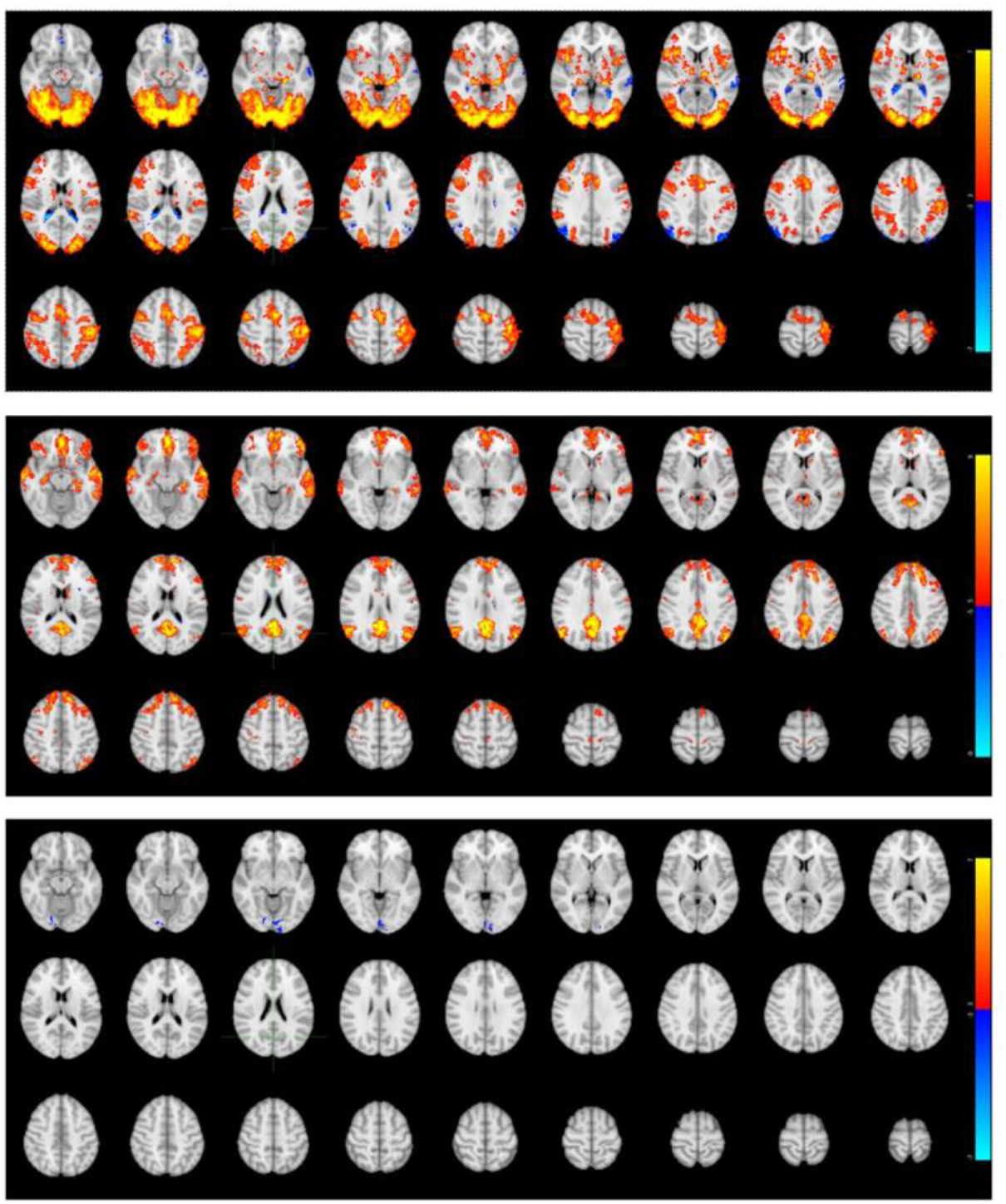
The volumetric version of the activation/deactivation patterns shown in Figures 2a, 2b, and 2c.

**S8:**
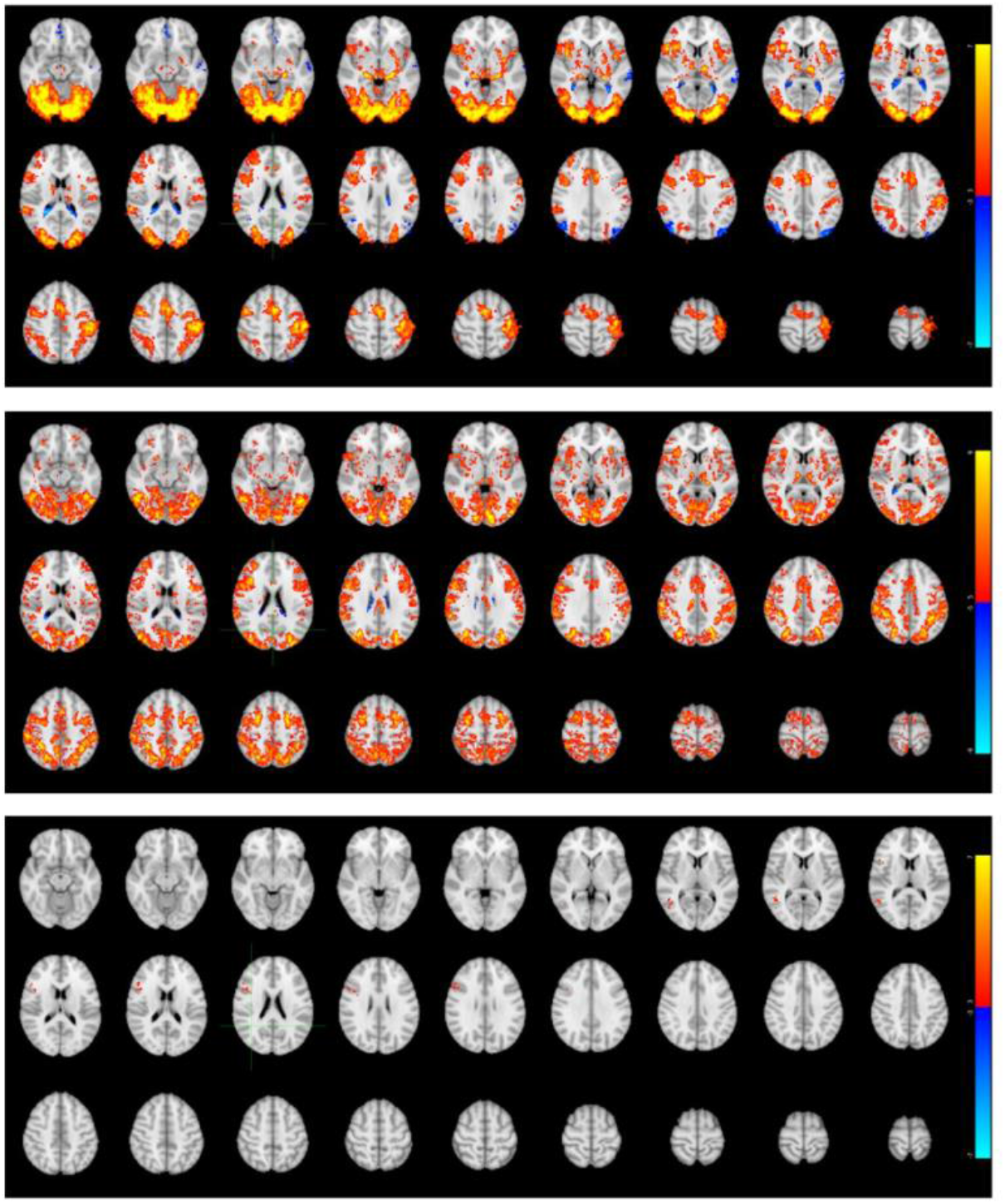
The volumetric version of the activation/deactivation patterns shown in Figures 2d, 2e, and 2f.

## Notes

### Competing Interest Statement

The authors have declared no competing interest.

